# Acute bee paralysis virus regulation of microRNA expression dynamics in the insect host (*Apis mellifera*) cell line, AmE-711

**DOI:** 10.1101/2025.09.19.677441

**Authors:** Deepak Kumar, Michael Goblirsch, John Adamczyk, Shahid Karim

## Abstract

**Background:** Honey bees (*Apis mellifera*) are essential pollinators that support global food production and economic stability. Their health and survival are threatened by diminishing floral resources, pesticide exposure, parasitic mites, and microbial and viral diseases. Among these stressors, viral infections are particularly challenging, often exacerbated by the parasitic mite, *Varroa destructor*, a competent vector of multiple honey bee viruses. Understanding the mechanisms underlying honey bee-virus interactions is critical for mitigating the negative impact of infections on colony health. One understudied aspect is the role of microRNAs (miRNAs) in viral pathogenesis and antiviral defense. miRNAs are short, non-coding RNAs produced by both hosts and pathogens that act as post-transcriptional regulators of gene expression and can influence host-pathogen dynamics during infection. In this study, we used a honey bee-derived cell line to test the hypothesis that viral infection perturbs honey bee- and viral-encoded miRNA expression.

**Methods:** Small RNA libraries from honey bee AmE-711 cells subjected to one of four treatments: media change only (uninfected), heat-killed Acute bee paralysis virus (ABPV), the viral mimic Poly(I:C), or infectious ABPV, were prepared using an Illumina Truseq kit. Sequencing data were analyzed using miRDeep2 and sRNAtoolbox to identify differentially expressed (DE) miRNAs, which were subsequently validated by RT-qPCR assay.

**Results:** Sequencing yielded > 3.6 x 10^8^ raw reads that were assigned to 12 small RNA libraries, from which, 481 unique miRNAs were identified. Moreover, 15 miRNAs were DE in ABPV-infected cells compared to uninfected cells: miR-2b-5p, miR-33-5p, miR-133-3p, miR-6001-3p, miR-996-3p, miR-965-3p, miR-125-5p, miR-13b-3p, miR-79-3p, miR-971-3p, miR-277-3p, miR-92c-5p, miR-6065-3p, miR-965-5p, and miR-3786-5p. We highlight some of the DE miRNAs identified in ABPV-infected cells that show regulatory effects in other systems in response to infection.

**Conclusion:** This study identified miRNAs differentially expressed in ABPV-infected cells, suggesting roles in either antiviral defense or in promoting viral pathogenesis through suppression of host immune responses. These results provide a foundation for functional studies using honey bee cell lines to clarify the cellular mechanisms governing honey bee-virus interactions.

## Introduction

Among the pathogens that infect honey bees (*Apis mellifera*), viruses are arguably the most challenging for beekeepers to control. Honey bee social living (Chen et al., 2006; Schmid-Hempel, 1995), beekeeper management of colonies (Alger et al., 2018; Owen, 2017; Fries and Camazine, 2001), pervasive infestation with the parasite and viral vector, *Varroa destructor* (Abban et al., 2024; Lin et al., 2022; Wilfert et al., 2016; Ryabov et al., 2014), and the absence of effective, widely available antiviral measures all facilitate transmission within and between honey bee colonies. Most disease-causing viruses identified to date in honey bees are positive-strand RNA (+ssRNA) viruses, and include representatives from Dicistroviridae (acute bee paralysis virus [ABPV], black queen cell virus, Israeli acute paralysis virus, and Kashmir bee virus), Iflaviridae (deformed wing virus [DWV], Kakugo virus, sacbrood virus, and slow bee paralysis virus), as well as unclassified viruses (chronic bee paralysis virus and Sinai viruses) (Grozinger and Flenniken, 2019).

Understanding the complexities of gene regulation, particularly in arthropod responses to infectious agents, has brought increasing attention to microRNAs (miRNAs). These short (21–25 nucleotides [nt]), single-strand non-coding RNAs are abundantly expressed by both hosts and pathogens during infection and can function by degrading host or pathogen-derived RNAs (Wang et al., 2022) or repressing protein translation through interaction with the RNA-induced silencing complex (Pasquinelli et al., 2000; Lee et al., 1993; Wightman et al., 1993). miRNA targeting of host or pathogen-derived RNAs is largely determined by complementary binding between the 3’-untranslated region of the RNA and the miRNA seed sequence, though multiple miRNAs with homologous seed sequences can share overlapping targets (O’Brien et al., 2018). In arthropods, miRNA-mediated regulation influences key cellular pathways involved in development, immunity, and pathogen resistance (Cao et al., 2024; Reynolds, 2024; Wei et al., 2018; Singh et al., 2012; Liu et al., 2009). Despite this, the functional role of miRNAs in honey bee innate immunity in response to viral infection remains largely unexplored.

Viral infections are widespread, and viral loads are abundant in honey bee colonies, making these pathogens a major factor in colony mortality (Lamas et al., 2025; Wilfert et al., 2016; Bowen-Walker et al., 1999). With no commercially available antiviral treatments, there is an urgent need to better understand honey bee-virus interactions.

Defining the molecular responses of honey bees to viral entry and replication is critical for developing strategies to mitigate unwanted colony mortality. Among the viruses that cause disease in honey bees, ABPV is frequently detected, often as a co-infection with other viruses or *V. destructor* infestation and is consistently linked to deteriorating colony health (Lamas et al., 2025; Francis et al., 2013; Berthoud et al., 2010; Genersch et al., 2010; Berenyi et al., 2006; Nordström et al., 1999).

In this study, we used small RNA sequencing to examine the miRNA-mediated regulatory landscape in ABPV-infected AmE-711 honey bee cells. Our objective was to identify novel and differentially expressed (DE) miRNAs that may contribute to the honey bee antiviral response. We detected 481 putative miRNAs in the AmE-711 cell culture system and found that 15 were significantly DE following ABPV infection. These DE miRNAs represent promising candidates for functional characterization and may reveal novel components of the honey bee innate immune defense. Given the central role of miRNAs in post-transcriptional gene regulation, these findings provide new insight into how honey bees modulate gene expression in response to viral pathogens.

## Materials and Methods

### AmE-711 Cells and ABPV Infection

AmE-711 honey bee cells were obtained by the USDA ARS through an agreement with the University of Minnesota (Minneapolis, MN, USA). AmE-711 is persistently infected with the +ssRNA virus, DWV (Carrillo-Tripp et al., 2016), a common pathogen of honey bee colonies worldwide (Wilfert et al., 2016). AmE-711 cells are routinely cultured in sealed 75 cm^2^ flasks (Corning, Arizona, USA) at 32°C in a non-humidified incubator.

The base growth medium is Schneider’s Insect Medium containing L-glutamine and sodium bicarbonate (Millipore Sigma, Burlington, MA), supplemented with 10% heat-inactivated fetal bovine serum (FBS; Millipore Sigma, Burlington, MA, USA).

For experiments, cells were detached from stock flasks using 0.25% trypsin-EDTA (Thermo Fisher Scientific, Waltham, MA, USA) and transferred to 6-well plates at ∼2.0 x 10^6^ cells per well. Two independent pools of cells were used to prepare two sets of 6- well plates per pool. Within each plate, sets of three wells were assigned to one of four treatments (Figure 1):

1) Medium change alone (uninfected control)
2) Exposure to heat-killed ABPV
3) Treatment with 20 µg/mL of double-stranded RNA analog Poly(I:C) (Millipore Sigma, Saint Louis, MO, USA) to simulate viral infection
4) Infection with viable ABPV

**Figure 1.**
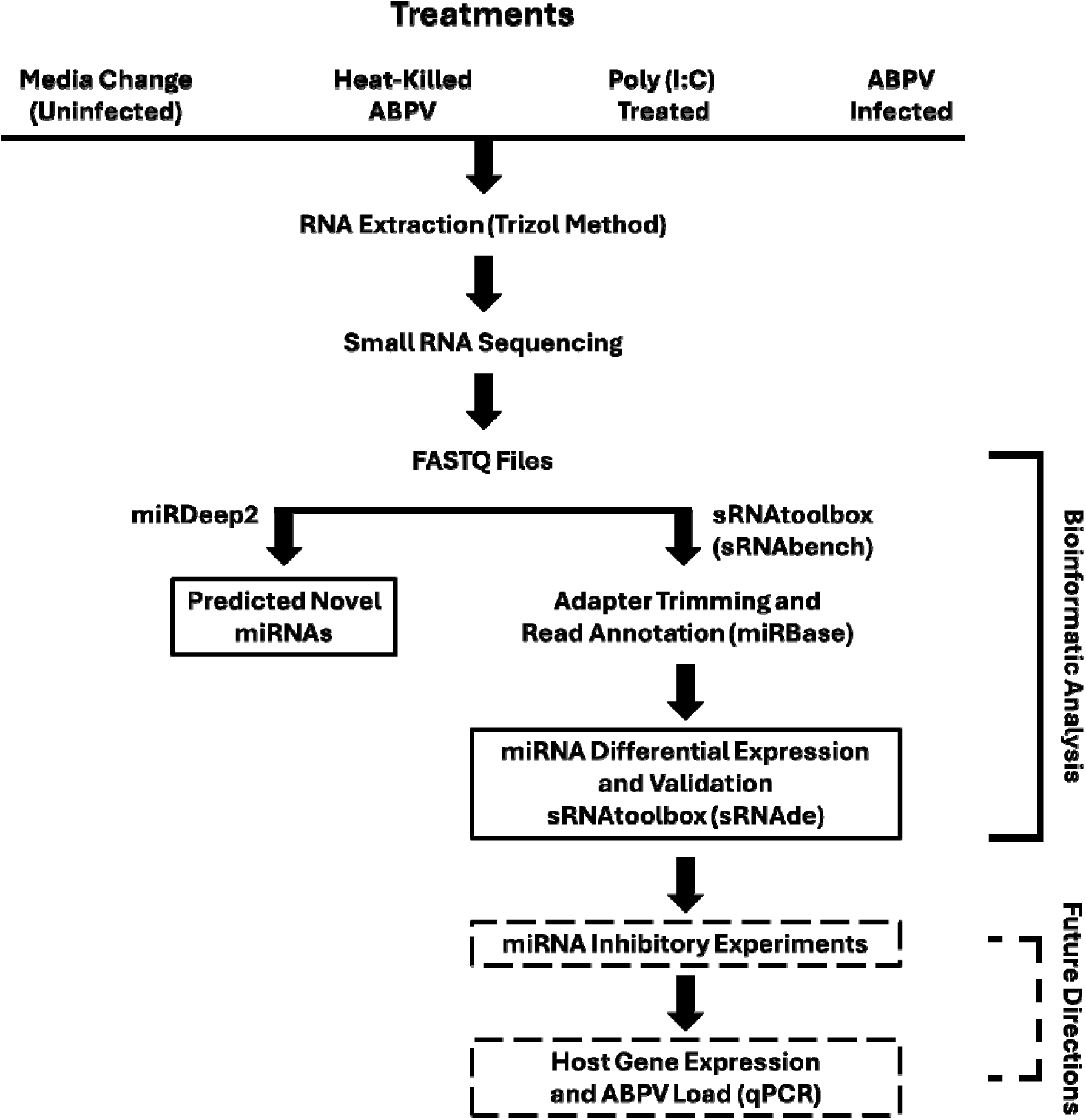
Graphical depiction of experimental workflow. AmE-711 honey bee cells were exposed for 24 hours to one of four treatments: media change alone as an uninfected control; heat-killed acute bee paralysis virus (ABPV); 20 µg/mL of double-stranded RNA analog Poly(I:C) to simulate viral infection; or infection with viable ABPV. Samples were then processed for small RNA sequencing. Sequence data were analyzed to identify differentially expressed (DE) miRNAs, which were subsequently validated by RT-qPCR assay. Future directions are highlighted with dashed boxes.

The ABPV inoculum originated from an AmE-711 culture that became contaminated with the virus. To propagate the virus, the medium from the contaminated culture was used to inoculate a flask containing healthy AmE-711 cells overnight. The infectious medium was then collected, passed through a 0.22 µm syringe filter, and stored as concentrated aliquots at -80°C. For heat-killed ABPV, 1 mL of the inoculum was heated to 95°C for 1 hour using a heat block and cooled to room temperature before dilution in Schneider’s Insect Medium. Infectious ABPV was similarly diluted to a concentration that produces observable cytopathic effects within 24 hours after inoculation while allowing recovery of sufficient high-quality RNA for downstream analysis. For each treatment, 1 mL of appropriate medium was added to the wells after a PBS wash. All treatment media lacked FBS. Cells were incubated with the treatment media for 24 h, after which the media were removed and 1 mL of TRIzol was added to each well. Lysates were collected and stored at -80°C until RNA was extracted.

### RNA Extraction, Library Preparation, and Sequencing

Total RNA was extracted from samples and RNA quality was assessed with a Nanodrop spectrophotometer. Small RNA libraries were generated at the Molecular and Genomics Core, University of Mississippi Medical Center (Jackson, MS, USA). Twelve libraries, four treatment groups with three biological replicates each, were prepared using the Illumina Revvity NextFlex V4 Small RNA kit (San Diego, CA, USA). Briefly, short adapter sequences were ligated to the small RNAs, followed by reverse transcription to cDNA and PCR amplification to incorporate sample-specific barcodes and sequencing adapters. Library concentrations were measured with a Qubit fluorometer, and fragment sizes were verified using an Agilent 2100 Bioanalyzer with a high-sensitivity DNA 100 chip after gel purification. Libraries were pooled and sequenced in single-read 51-base mode on the Illumina NextSeq 2000 platform using the NextSeq 2000 P3 Reagents kit (San Diego, CA, USA).

### Small RNA Data Processing and Analysis

Small RNA datasets were analyzed using the sRNAtoolbox web server (https://arn.ugr.es/srnatoolbox/) as described by Aparicio-Puerta et al. (2022), with the honey bee reference genome GCA_003254395.2_Amel_HAv3.1_genomic. Sequence quality was first assessed using FastQC (http://www.bioinformatics.babraham.ac.uk/projects/fastqc). Preprocessing, mapping, and annotation were performed primarily with the sRNAbench module (Aparicio-Puerta et al., 2022), supplemented with custom scripts when needed. Adapter detection and trimming followed an iterative approach. Adapters were first searched across the full read length, and if not detected, progressively shorter 3’-end segments were examined. After adapter removal, reads collapsed into unique sequences, and read counts were assigned to quantify sequence abundance. As sRNAtoolbox has limitations in the discovery and annotation of novel miRNAs, miRDeep2 (V2.0.0.8; Friedländer et al., 2012) was also used to predict both novel and known miRNAs from the datasets.

### Differential Expression Analysis

Differentially expressed miRNAs were identified using the sRNAde module (Robinson et al. 2010). An expression matrix of raw read counts was generated and analyzed with edgeR, applying the trimmed mean of M-values normalization method (Robinson et al., 2010). In addition, sRNAbench was used to generate an expression matrix normalized to reads per million using the single-assignment method, in which reads mapping to multiple miRNAs were assigned only to the miRNA with the highest expression level.

This method primarily affected reads aligning to multiple reference sequences within the same miRNA family. Reads per million values were calculated by dividing the read count for each miRNA by the total number of reads mapped to the miRNA library.

### miRNA Target Prediction and Functional Analysis

The miRNAconsTarget tool within the sRNAtoolbox was used to predict honey bee genes potentially regulated by DE miRNAs. This tool integrates three targeting algorithms: TargetSpy (Sturm et al., 2010), MIRANDA (John et al., 2005), and PITA (Kertesz et al., 2007). Genes predicted as targets by all three algorithms were prioritized for further analysis. Although *in silico* target prediction can yield false positives, refinement through cross-species comparisons and consideration of combinatorial targeting reduced inaccuracies (Min and Yoon, 2010). Predicted target genes were functionally characterized using the STRING (Franceschini et al., 2013) and PANNZER2 (Törönen et al., 2018) web servers. STRING provided gene interaction networks and identified significantly enriched pathways, while PANNZER2 was used to reannotate predicted protein targets. Gene ontology (GO) annotation analysis and visualization were performed with SRplot (Tang et al., 2023).

### RT-qPCR Validation of Differentially Expressed miRNAs

Differentially expressed miRNAs were validated using RT-qPCR. cDNA synthesis and miRNA expression profiling were performed with the Mir-X miRNA RT-qPCR TB Green kit (Takara BIO, San Jose, CA, USA). Total RNA was first polyadenylated using poly(A) polymerase and then reverse transcribed using SMART MMLV reverse transcriptase.

The resulting cDNA was amplified using the TB Green Advantage qPCR Premix, together with the mRQ 3’ primer and miRNA-specific 5’ primers. The RT-qPCR cycling protocol included an initial denaturation step of 95°C for 10 minutes, followed by 40 cycles of 95°C for 5 seconds and 60°C for 20 seconds.

## Results and Discussion

### Length Distribution and Biotype of Small RNAs

Small RNA sequencing generated > 3.6 x 10^8^ raw reads from 12 libraries. After adapter trimming and removal of reads < 20 nt and > 30 nt, > 2.4 x 10^8^ reads remained. The types of small RNAs present in the samples are reflected in the distribution of read length (Figure 2). The distribution showed two prominent peaks, one at 22 nt, corresponding to miRNAs and small interfering RNAs (siRNAs), and another between 26 and 30 nt, indicating PIWI-interacting RNAs (piRNAs). The presence of piRNAs in honey bees has been confirmed in other studies (Sun et al., 2023; Watson et al., 2022; Wang et al., 2017). piRNAs associate with PIWI proteins, members of the Argonaute family, and play key roles in germline and stem cell development in invertebrates. These small RNAs are essential for silencing transposable elements, thereby preserving genome integrity (Aravin et al., 2008; Brennecke et al., 2008; Brennecke et al., 2007).

**Figure 2.**
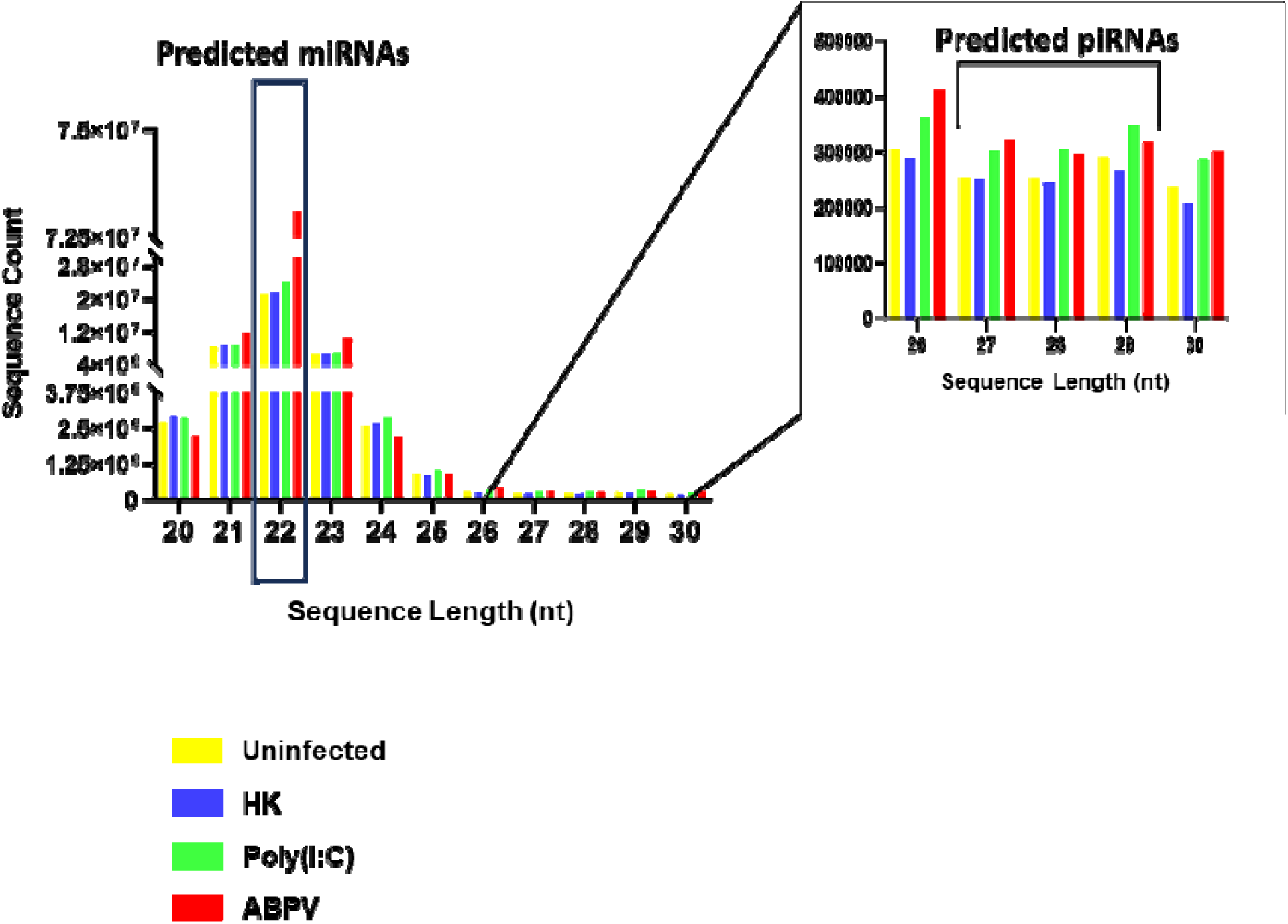
Distribution of > 2.4 x 10^8^ small RNA sequence reads 20–30 nucleotides (nt) in length generated from AmE-711 cells exposed to media change only (uninfected; yellow), heat-killed (HK) acute bee paralysis virus (ABPV) (blue), Poly(I:C) (green), and infectious ABPV (red). The main panel (left) emphasizes the frequency of microRNAs (22 nt) while the inset (right) highlights the distribution of piRNAs (26–30 nt).

In mosquitoes, piRNAs also target viruses, including arboviruses (Leger et al., 2013; Schnettler et al., 2013; Morazzani et al., 2012; Vodovar et al., 2012; Hess et al., 2011) and insect-specific viruses, such as cell-fusing agent virus and Phasi Charoen-like virus (Whitfield et al., 2017). Beyond mosquitoes, demonstration of piRNA antiviral activity is limited; *Diaphorina citri* remains the only non-mosquito insect where viral infection by densovirus triggers a ping-pong piRNA response (Nigg et al., 2020). Functional evidence supports the antiviral role of piRNAs in mosquitoes where RNAi knockdown of PIWI proteins 4-6 in *Aedes aegypti* reduces piRNA production and increases viral replication (Tassetto et al., 2019; Dietrich et al., 2017; Whitfield et al., 2017; Varjak et al., 2017).

In *A. mellifera*, the best characterized and primary antiviral defense mechanism is the siRNA pathway (Brutscher and Flenniken, 2015; Vijayendran et al., 2013; Ding, 2010; Hammond et al., 2001). One likely reason for mosquitoes to invest more heavily in piRNAs for antiviral defense is their expanded encoding of PIWI genes. While *A. mellifera*, *Bombyx mori*, and *Drosophila* sp. encode up to three PIWI genes, *Ae. aegypti* encodes seven (Lewis et al., 2018), potentially broadening its piRNA target range. Future work will focus on identifying and characterizing piRNAs in ABPV-infected cells to determine whether they play a direct role in honey bee antiviral defense.

### Other Small RNA Categories

Additional small RNA categories included miRBase (sense), messenger (sense and antisense; mRNA), noncoding (ncRNA), ribosomal (rRNA), small nuclear (snRNA), small nucleolar (snoRNA), transfer (tRNA), and unassigned and other RNAs (Figure 3). miRBase (sense) sequences, which represent the nucleotide composition of mature miRNAs transcribed from a DNA template, were the most prevalent, regardless of treatment. In ABPV-infected cells, miRBase (sense) miRNAs comprised 53.89% compared to 70.25%, 71.97%, and 73.11% for the Poly(I:C), uninfected, and heat-killed ABPV treatments. The reduction in miRbase (sense) miRNAs in ABPV-infected cells could suggest a countermeasure of the virus to suppress honey bee miRNA biogenesis (Han et al., 2025), shifting the small RNA pool towards virus-derived small RNAs or host-derived RNA degradation products. Notably, ABPV-infected cells had a markedly higher proportion of unassigned reads (22.73%) compared to cells from the heat-killed ABPV (8.10%), uninfected (8.36%), and Poly(I:C) (8.91%) treatments. This pattern is typical for virus-infected samples, where novel or fragmented small RNAs are abundant (Aguiar et al., 2016). Elevated rRNA (9.65%) and sense and antisense mRNA fragments (7.48%) in ABPV-infected cells may further reflect virus-induced host RNA degradation. In contrast, other RNA categories including ncRNA, snRNA, snoRNA and tRNA remained low, each < 1%, and stable across treatments, likely representing baseline cellular RNA turnover. Interestingly, heat-killed ABPV and Poly(I:C) treatments did not induce shifts comparable to those seen in ABPV-infected cells, indicating that these changes in miRNA abundance and read composition are specific to active ABPV infection and replication rather than general immune activation.

**Figure 3.**
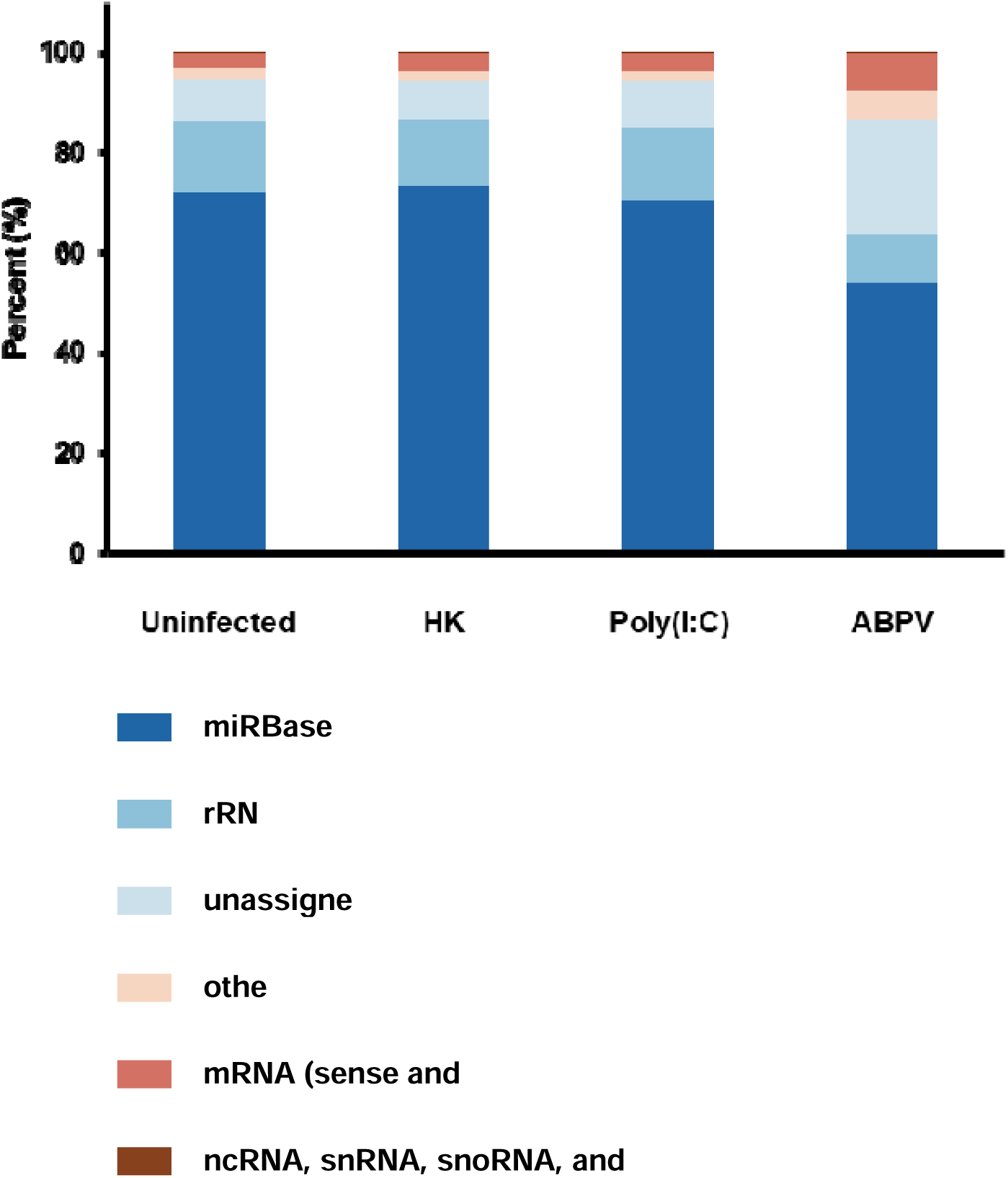
The proportional abundance of different categories of small RNA reads in AmE-711 cells exposed to each of the four treatment conditions: media change only (Uninfected), heat-killed (HK) acute bee paralysis virus (ABPV), Poly(I:C)-treated, and ABPV-infected.

### *In Silico* Mapping of Small RNA Sequences to the ABPV Genome

We aligned small RNA sequences from ABPV-infected cells to the ABPV reference genome (GCA_000856345.1_ViralProj14983_genomic). In infected cells, 16.5% of the sequences mapped to the ABPV genome. In contrast, only 0.022% and 0.016% of the sequences from heat-killed ABPV and uninfected cells, respectively, aligned to the viral genome, indicating successful infection in cells inoculated with virus and effective heat inactivation and minimal background contamination in the respective controls. The high proportion of viral-aligned reads in infected cells reflects active ABPV replication and underscores the susceptibility of AmE-711 cells to infection. These findings support the use of the AmE-711 cell line as a model for investigating ABPV pathogenesis, host-virus interactions, and antiviral responses.

### Comparison of miRNA Expression Between ABPV-Infected and Uninfected Cells

We identified 481 unique miRNAs, including 142 that are honey bee-specific and 270 that are orthologs to miRNAs from other organisms cataloged in miRBase. We also predicted 69 novel miRNAs; however, applying a miRDeep2 score threshold of < 4 (Hermance et al., 2019), reduced this number to 46 (Supplementary Data link - <u>Supplementary data .xls</u>).

Fifteen miRNAs were DE in ABPV-infected cells compared to uninfected cells: miR-2b- 5p, miR-33-5p, miR-133-3p, miR-6001-3p, miR-996-3p, miR-965-3p, miR-125-5p, miR-13b-3p, miR-79-3p, miR-971-3p, miR-277-3p, miR-92c-5p, miR-6065-3p, miR-965-5p, and miR-3786-5p (Figure 4A). Some of the known functions of the DE miRNAs are associated with cellular regulation during viral infection (Table 1). For example, miR-2b- 5p was significantly upregulated in ABPV-infected cells compared to uninfected cells. In *Ae. aegypti* Aag2 cells infected with chikungunya virus (CHIKV), miR-2b-5p was also upregulated and shown to target ubiquitin-related modifier transcripts, ultimately leading to the suppression of viral replication (Dubey et al., 2017). Conversely, the opposite effect was observed in *Ae. albopictus* and *Anopheles stephensi* cells infected with CHIKV and *Plasmodium* sp., respectively, where miR-2b was downregulated and did not target immune signaling transcripts in response to infection (Shrinet et al., 2014). Given this evidence, the upregulation of miR-2b-5p in ABPV-infected cells may reflect an evolutionarily conserved antiviral mechanism. Its increased expression in ABPV- infected cells may contribute to restricting replication by targeting viral transcripts. Further work is needed to identify its specific targets in *A. mellifera* and to confirm its functional role in honey bee antiviral immunity.

**Figure 4.**
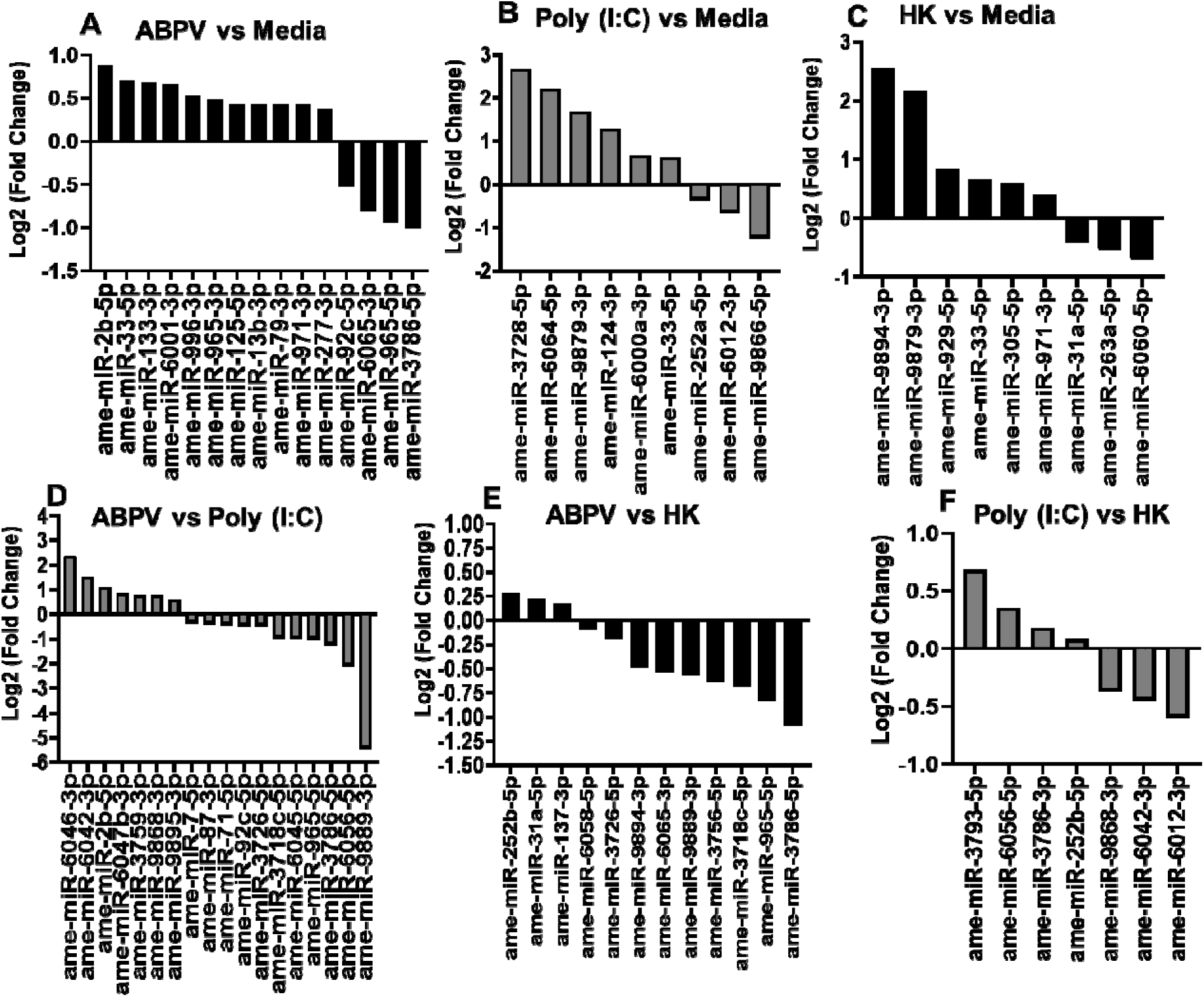
*In silico* differential expression (DE) of microRNAs (miRNAs) in AmE-711 cells. A) acute bee paralysis virus-infected (ABPV) v. uninfected cells (media); B) Poly(I:C)-treated v. uninfected (media); C) Heat-killed (HK) ABPV exposure v. uninfected cells (media); D) ABPV vs Poly (I:C); E) ABPV v. HK; F) Poly(I:C) vs HK. EdgeR was used for DE analysis. All DE miRNAs are statistically significant (*p* value ≤ 0.05) relative to the second group mentioned in each panel.

**Table 1.** Differentially expressed microRNAs (miRNA) detected in AmE-711 cells infected with acute bee paralysis virus compared to uninfected cells, their putative role(s), and gene target(s).

| miRNA | Putative function | Putative gene target | Reference |
| --- | --- | --- | --- |
| 2b-5p | Chikungunya virus replication in *Aedes aegypti* cells | *ubiquitin-related modifier* | Dubey et al., 2017 |
| 13b-3p | Interaction with Imd pathway to negatively regulate response to Gram-negative bacteria; slow bee paralysis virus pathogenesis | Insulin-like growth factor 2 mRNA-binding protein 1-like (LOC100647990); ecdysone-induced protein 75B, isoforms C/D-like, (LOC100644185) | Wei et al., 2018; Li et al., 2017; Niu et al., 2017 |
| 33-5p | Dengue type 2 virus pathogenesis |  | Campbell et al., 2014 |
| 79-3p | Innate immunity; differentiation; apoptosis | Roundabout protein 2 pathway (robo2), DRAPER, HP2 (Hemolymph protease), P38, pvr, puc | Artigas-Jerónimo et al., 2019; Dong et al., 2017; Ouyang et al., 2015; Seddiki et al., 2013; Pedersen et al., 2013; Fullaondo and Lee., 2012; Yuva-Aydemir et al., 2011; Bond and Foley, 2009 |
| 92c-5p | Adult eclosion; honey bee neural resistance to fluvalinate exposure |  | Li et al., 2023; Xueqing et al., 2023 |
| 125-5p | aae-miR-125-5p/hsa-miR-125b-5p target human mRNAs involved in immune and inflammatory responses; viral replication in mosquitoes | *RIG-I* (retinoic acid-inducible gene I), *DHX58* (DEXH box polypeptide 58), and *TIRAP* (toll/interleukin-1) | Fiorillo et al., 2022; Wu et al., 2020; Maharaj et al., 2015 |
| 133-3p | Hemocyte proliferation and innate immunity of *Scylla paramamosain*; Tick-borne *Anaplasma phagocytophilum* survival and transmission | *astakine*, *isoatp4056* | Zhang et al., 2022; Ramaswamy et al., 2020 |
| 277-3p | Caste differentiation in the bumblebee, *Bombus lantschouensis* |  | Liu et al., 2019 |
| 965-3p | White spot syndrome virus infection and shrimp antiviral immunity mediated by targeting the virus gene *wsv240* and the host phagocytosis-related gene *ATG5* | *gamma-interferon-inducible lysosomal thiol reductase, cytohesin-1, Alk, GTP/GDP exchange factor, MLK1, STAT, TEP1* | Li et al., 2017; Shu et al., 2016; Zhang et al., 2014 |
| 971-3p | Stimulation of female reproduction in corpora allata; development (conserved across all insects), chemosensory signaling in honeybee | Juvenile hormone synthesis pathway genes, *wat* (involved in chemosensory signaling) | Li et al., 2025; Ma et al. 2021; Shi et al., 2017 |
| 996-3p | Begomovirus infection and transmission in whitefly, *Bemisia tabaci*; Regulation of vitellogenin in the ectoparasitic caligid copepod, *Caligus rogercresseyi* | *bta08371* | Casuso et al., 2024; Hasegawa et al., 2020 |
| 6001-3p | Developmental processes of *Apis mellifera ligustica* worker's midgut; development of larval gut of *Apis cerana* |  | Fan et al., 2023; Guo et al., 2018 |
| 6065-3p | Linked to eusocial Aculeata |  | Sovik et al., 2015 |

Another miRNA, miR-33-5p, was upregulated in ABPV-infected cells compared to uninfected cells. This finding is not unique to invertebrates, as differential expression of miR-33-5p has been observed in various cells or tissues of chickens and ducks infected with virus (Yang et al. 2020; Li et al., 2015) and *Mycoplasma gallisepticum* (Sun et al., 2022). However, it does contrast with previous reports in mosquitoes, where miR-33-5p was downregulated in response to infection with Dengue virus (Campbell et al., 2014). Su et al. (2017) also observed downregulation of miR-33-5p in mosquitoes given a virus-free bloodmeal; however, several miRNAs, including miR-1175, miR-276, and miR-371, displayed opposite expression patterns in *Ae. aegypti* and *Ae. albopictus*, highlighting that the regulatory roles of miRNAs can vary significantly across species. These observations emphasize the importance of the species-specific context in miRNA-mediated antiviral responses. For example, in *Drosophila* sp., miR-33a and –b have roles other than in antiviral immunity by targeting transcripts that regulate fatty acid metabolism and insulin signaling (Dávalos et al., 2011). Further mechanistic studies are needed to determine the functional role of miR-33-5p in honey bee antiviral immunity.

Our data suggest that upregulation of miR-133 in ABPV-infected cells may represent a honey bee innate immune response aimed at suppressing viral proliferation. Evidence from other invertebrate models supports this interpretation. In the mud crab, *Scylla paramamosain*, miR-133 was upregulated following infection with either white spot syndrome virus (WSSV) or *Vibrio parahaemolyticus*, and inhibition of miR-133 using anti-miRNA led to increased WSSV levels (Zhang et al., 2022). Similarly, Ramaswamy et al. (2020) showed that miR-133 targets and inhibits organic anion transporting polypeptide *isoatp4056* expression in *Ixodes scapularis*, and that treatment with miR- 133 mimic or miR-133 precursor reduced *Rickettsia* sp. load in infected ticks, with potential implications for pathogen transmission. Another potential target of miR-133 is *astakine*, a hematopoietic factor and immune regulator (Zhou et al., 2023). Adegoke et al. (2023) reported a three-fold increase in *AmHem-345670* transcript, a coding sequence for *astakine*, in response to *Rickettsia parkeri* infection in ticks, indicating a potential role for *astakine* in facilitating *R. parkeri* replication and proliferation. Zhang et al. (2022) further showed that miR-133 knockdown in virus- or bacteria-infected mud crabs led to elevated *astakine* expression, increased hematopoiesis and apoptosis, and reduced survival. Taken together, these studies indicate that miR-133 plays an important role in immune regulation across arthropods and that its upregulation in ABPV-infected cells points to a potential inhibitory function in honey bee viral infections.

One miRNA found to be upregulated in our study, miR-6001, was previously identified as a factor involved in immature development of the Asian honey bee, *A. cerana*. Fan et al. (2023) reported constitutive expression of miR-6001-y in the larval gut of *A. cerana*, and through GO and KEGG analyses, connected it to key developmental signaling pathways, including Wnt, Hippo, and Notch. In terms of viral pathogenesis, Wnt signaling has been shown to be negatively correlated with Rift Valley fever virus infection in *Ae. aegypti* (Smith et al., 2023). Similarly, this pathway exerts a strong antiviral effect against SARS-CoV-2 and other pathogenic RNA viruses *in vitro*, significantly reducing viral replication and load, inflammation, and clinical symptoms in a murine model of COVID-19 infection (Xu et al., 2024). The Wnt pathway is an evolutionarily conserved signaling cascade that regulates cellular proliferation, development, and self-renewal across many species. Importantly, several studies have shown that viruses can manipulate this pathway to enhance infection and replication (Harmon et al., 2016; Liu et al., 2011; Cha et al., 2004). Given that miR-6001 remains a relatively understudied miRNA, further research is needed to clarify its regulatory role in modulating this conserved pathway, particularly in the context of its potential antiviral function and precise mechanism of action in honey bee cell biology.

An increase in the expression of miR-996 in ABPV-infected cells was not surprising. Hasegawa et al. (2020) showed an increase in miR-996 expression in a whitefly model of tomato yellow leaf curl virus infection. The precise mechanism of miR-996 has yet to be investigated across different viral infection models of invertebrates. However, miR- 996 has been implicated in regulating vitellogenin in the caligid copepod (*Caligus rogercresseyi*), an ectoparasite of economic impact in the Chilean salmon industry (Casuso et al., 2024). The egg yolk precursor, vitellogenin, is a multifunctional protein recognized as a critical player in biological processes of invertebrates including reproduction, embryonic development, and immune responses. In the honeybee, vitellogenin is a well-studied hemolymph protein for that interacts with juvenile hormone in regulating adult worker behavioral development and lifespan via the nutrient signaling pathway (Amdam et al., 2004; Guidugli et al., 2005; Nelson et al., 2007). The interaction of vitellogenin and juvenile hormone is sensitive to disease conditions, it could lead to precocious foraging and premature death in workers infected with pathogenic microsporidia, viruses, and other stresses like agrichemical exposure and poor nutrition (Antúnez et al., 2009; Castelli et al., 2021; Coulon et al., 2020; Goblirsch et al., 2013; Lourenço et al., 2021).

Similar to what we observed in ABPV-infected cells, increased expression of miR-965 has been reported in other invertebrate host-virus interactions. For example, miR-965 levels increased in the Chinese white shrimp, *Fenneropenaeus chinensis*, after infection with WSSV (Li et al., 2017) and the diamondback moth, *Plutella xylostella*, infected with *Metarhizium anisopliae* (Zhang et al., 2024). One mechanism by which miR-965 functions in immune defense comes from a study on *Manduca sexta* larvae. Zhang et al. (2014) observed upregulation of miR-965 in hemocytes of bacteria-challenged moth larvae and that it targets genes in the JAK-STAT pathway, a well-known regulator of the host immune response. Another study demonstrated that miR-965 from the shrimp, *Marsupenaeus japonicus*, has antiviral activity against WSSV through two complementary mechanisms. First, miR-965 targets and suppresses the WSSV gene, *wsv240*, which is essential for infection (Shu et al., 2016). Second, miR-965 enhances the host’s innate immunity by targeting *autophagy 5*, an inhibitor of phagocytosis.

Suppression of *autophagy 5* leads to increased phagocytic activity of shrimp hemocytes (Shu et al., 2016). Together, these findings suggest miR-965 plays a dual role in antiviral defense by both directly suppressing viral genes and boosting host immune responses. Further research on honey bee-virus interactions is needed to identify the targets of miR-965, whether the miRNA is host- or virus-derived, and to understand how its inhibition or overexpression affects infection dynamics.

Interestingly, increased expression of miR-125-5p in ABPV-infected cells mirrors the response seen in *Ae. aegypti* and *Ae. albopictus* mosquitoes infected with CHIKV (Maharaj et al., 2015). Maharaj et al. (2015) suggested that mir-125-5p may play a role in regulating viral replication. Based on this, it is reasonable to speculate that miR-125- 5p could similarly influence ABPV replication in honey bee cells, although further research is needed to confirm this. In contrast, a different viral infection study found miR-125 was downregulated in ducklings infected with duck hepatitis A virus type 3 (DHAV-3), a significant pathogen in the duck industry that causes high mortality (Yugo et al., 2016). miR-125-5p has been shown to target retinoic acid-inducible gene I (*RIG-1*), DEXH box polypeptide 58, and toll/interleukin-1 receptor domain-containing adapter protein (Wu et al., 2020). When miR-125-5p is downregulated, *RIG-*I expression increases, which has been linked to enhanced DHAV-3 replication. *RIG-I* is a cytosolic pattern recognition receptor that detects viral RNA and triggers an antiviral response in many cell types (Chan and Gack, 2015).

Relative to uninfected AmE-711 cells, miR-6001-3p and miR-996-3p were found exclusively in ABPV-infected cells (Figure 4A), while miR-3728-5p, miR-6064-5p, and miR-124-3p were uniquely detected in cells treated with Poly(I:C) (Figure 4B). This difference in DE miRNAs likely reflects how AmE-711 cells respond to the distinct properties of these immunogenic stimuli. Poly(I:C) is a synthetic double-stranded RNA (dsRNA) that mimics viral infection, simulating dsRNA viruses (Fortier et al., 2004).

However, Poly(I:C) lacks capsid proteins that normally encase the viral genome in mature virions and serve as recognition factors for host cell receptors (Boulant et al., 2015). Despite these differences, some common DE miRNAs were observed between ABPV-infected and Poly(I:C)-treated cells. For example, miR-33-5p was upregulated in both treatment groups, suggesting host cell recognition of endogenous dsRNA. Future studies could explore changes in the expression of genes in the RNAi pathway to determine if Poly(I:C) triggers a response similar to ABPV.

Heat-killed ABPV should ideally have minimal impact on miRNA expression, as the virus can no longer establish infection (Elveborg et al., 2022). However, non-RNA viral components, such as proteins or structural elements may still elicit immune responses or other biological effects (Heilmann et al., 2025). Therefore, the DE miRNAs observed in cells exposed to heat-killed ABPV (Figure 4C) might result from these viral or other non-viral factors. Interestingly, two DE miRNAs, miR-965-5p and miR-3786-5p were shared between ABPV-infected cells and those exposed to heat-killed ABPV, which may suggest a lack of specificity in the immune response to foreign insult.

### Prediction of Target Genes, Gene Ontology, and Functional Enrichment Analyses of Target Networks

Honey bee proteins predicted to be targets of DE miRNAs were used to build a high-confidence interaction network (interaction score > 0.9; Figure 5). STRING analysis revealed that target proteins of 11 upregulated and 4 downregulated miRNAs exhibited significantly more interactions among themselves compared to a randomly selected set of *A. mellifera* proteins of similar size and degree distribution (nodes = 377, edges = 52, average node degree = 0.276, average local clustering coefficient = 0.109, expected number of edges = 25, PPI enrichment *p* value < 0.001). This enrichment analysis shows significant protein-protein interactions among the predicted targets of the DE miRNAs, strongly suggesting that these proteins are interrelated as a group and potentially participating in common biological processes or pathways.

**Figure 5.**
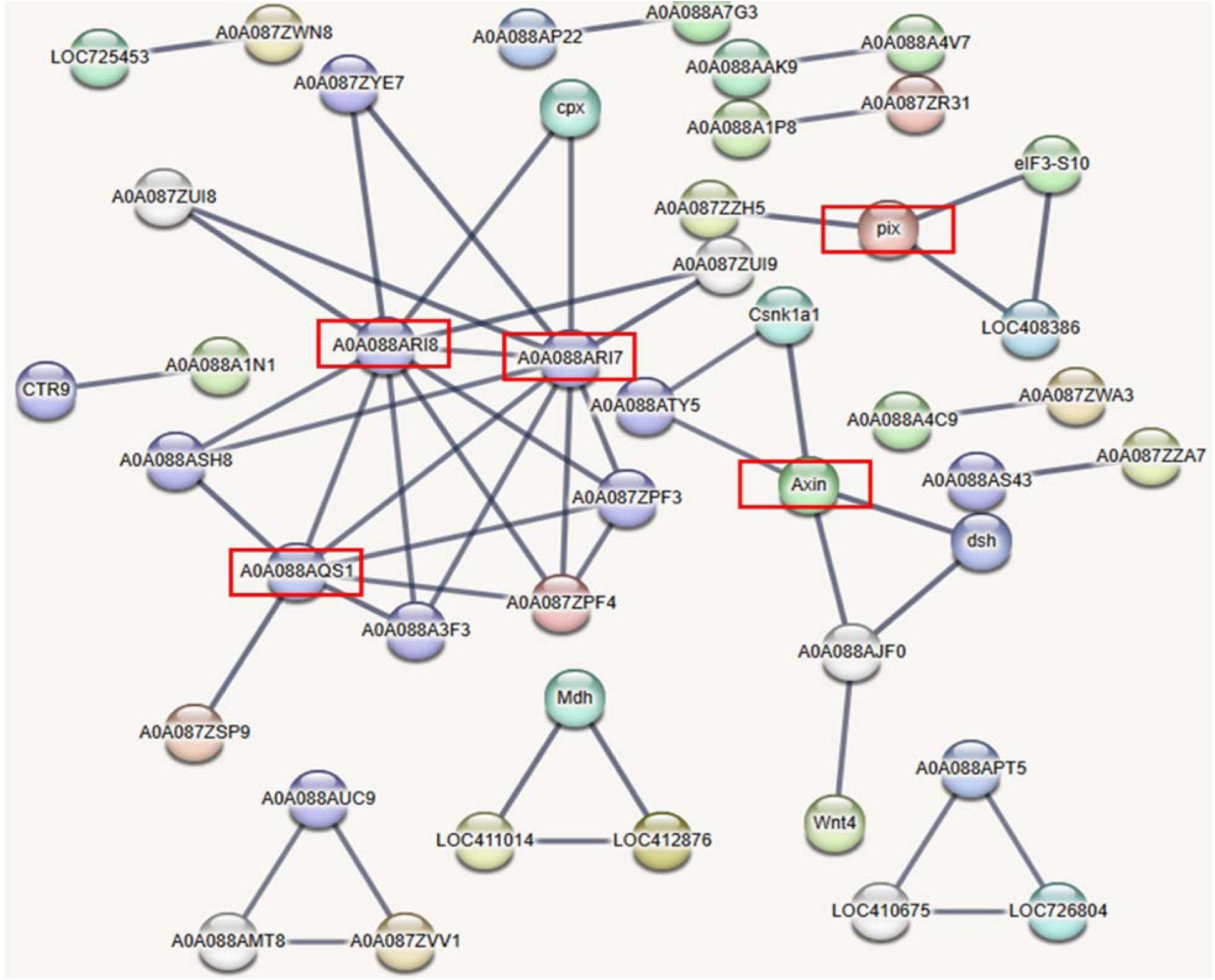
Network constructed exclusively from *Apis mellifera* proteins targeted by computationally identified differentially expressed miRNAs in cells infected with acute bee paralysis virus compared to uninfected cells. Red boxes highlight proteins involved in processes or pathways that may function in viral pathogenesis. These proteins include members of the Wnt signaling pathway (XP_006564443.2), which is known to be activated by RNA viruses, and proteins associated with exocytosis (XP_016769840.2 and XP_006564520.1), translational control (XP_006569310.1), and protein kinase activity (XP_006559979.1).

Numerous target genes corresponding to the DE miRNAs were predicted using the sRNAtoolbox miRNAconsTarget program (Aparicio-Puerta et al., 2019) (Figure 6). To minimize false positives, only targets predicted by all three miRNA target-prediction algorithms, TargetSpy, MIRANDA, and PITA, were considered. Gene ontology analysis showed that most target genes play important roles in cellular processes, metabolic processes, biological regulation, developmental processes, and responses to stimuli.

**Figure 6.**
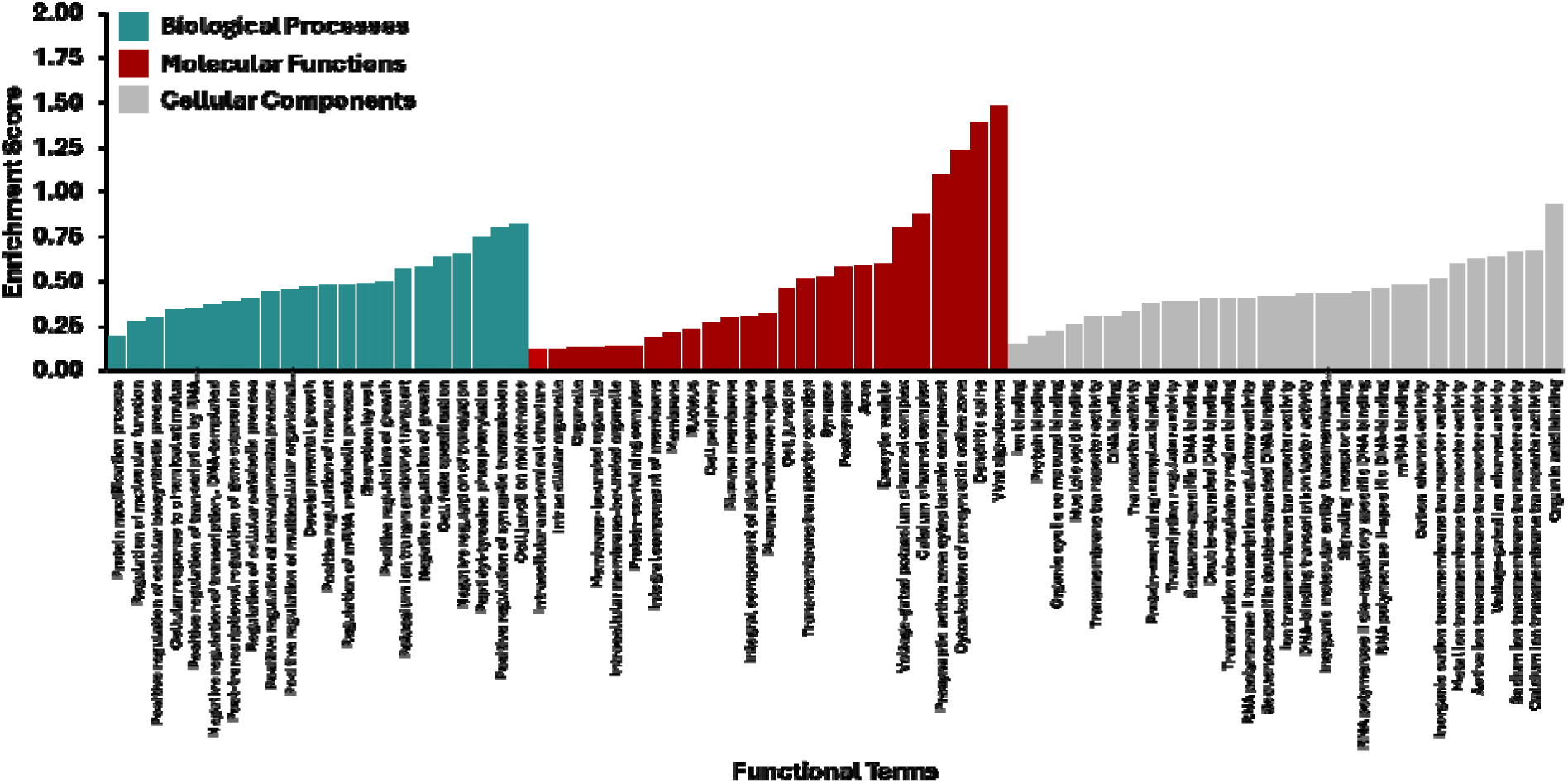
Gene ontology derived biological processes associated with *Apis mellifera* genes targeted by differentially expressed miRNAs in AmE-711 cells infected with acute bee paralysis virus compared to uninfected cells.

Pathway analysis using STRING and PANNZER2 highlighted involvement in Wnt signaling, exocytosis, translational control, and protein kinase activity, which are expected to be regulated by DE miRNAs in ABPV-infected cells. A detailed understanding of these miRNAs will be essential to clarify their roles in viral infection and the pathways they influence.

### Validation of *In Silico* Differentially Expressed microRNAs by RT-qPCR

The expression levels of DE miRNAs in ABPV-infected cells were evaluated using RT-qPCR (Figure. 7). For most of the miRNAs evaluated, the RT-qPCR results were consistent with those obtained from small RNA sequencing. Some discrepancies were observed between the two methods. These differences may stem from the distinct techniques used for miRNA quantification (Saldaña et al., 2017).

**Figure 7.**
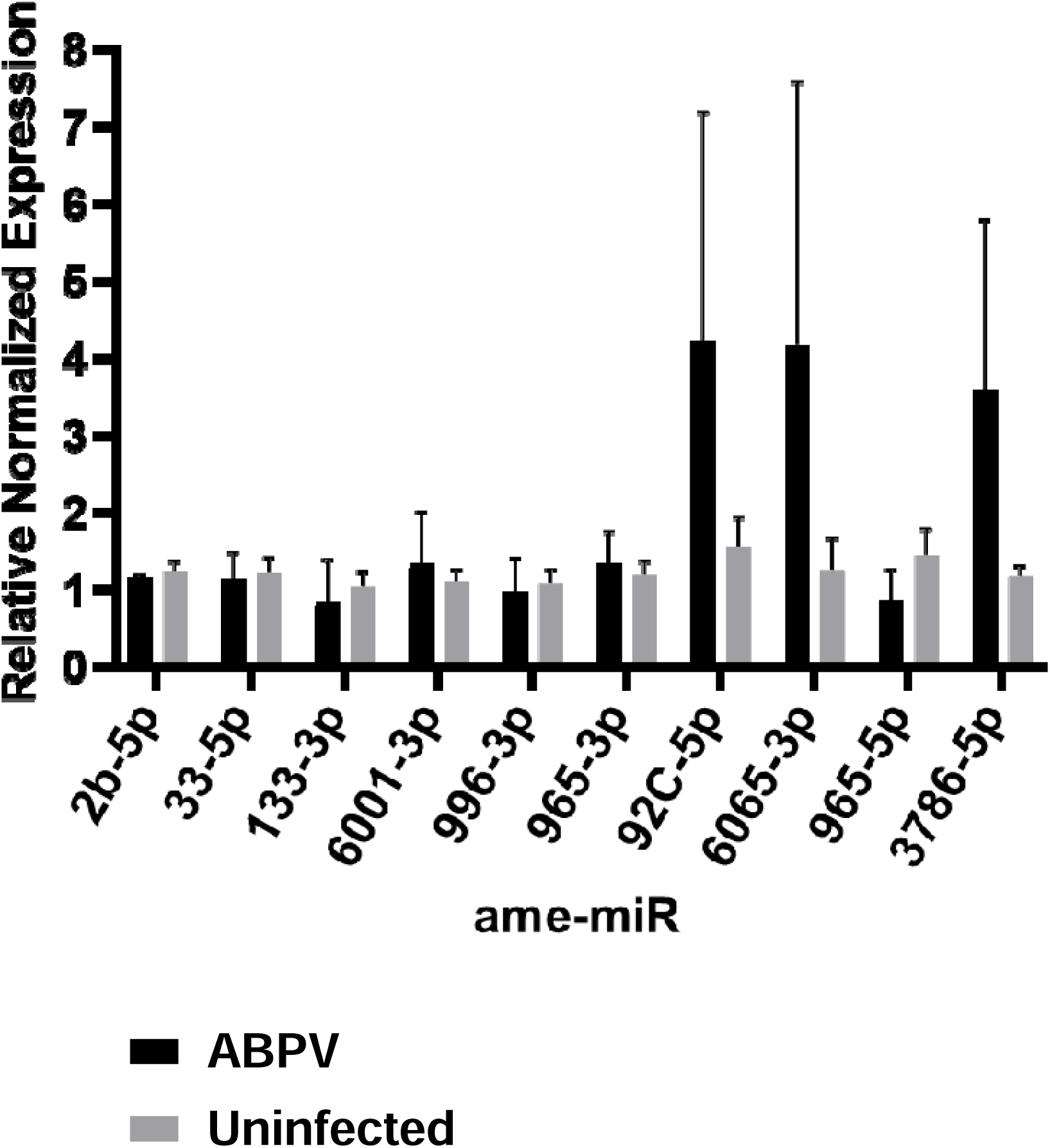
RT-qPCR validation of differentially expressed miRNAs (ame-miR) in cells infected with acute bee paralysis virus (ABPV) compared to uninfected cells.

## Conclusion

Our study established a miRNA profile of honey bee viral infection using host-specific cells infected with the +ssRNA virus, ABPV. This approach also provides a novel model for investigating viral infection mechanisms and pathogenesis *in vitro*, as well as for identifying molecular responses to relevant honey bee stressors at the cellular level. We identified several DE miRNAs in ABPV-infected cells. Notably, two miRNAs, miR-965-5p and miR-3786-5p, were DE in ABPV-infected cells compared to uninfected cells. Further investigation is needed to clarify the specific roles of these DE miRNAs in viral pathogenesis and the honey bee antiviral immune response.

Additionally, miR-33-5p was found to be upregulated in cells that were either infected with ABPV or treated with the dsRNA virus simulator, Poly(I:C), suggesting a potential role for this miRNA in host interactions with +ssRNA and dsRNA viruses. Other DE miRNAs, miR-3728-5p, miR-6064-5p, and miR-124-3p, were uniquely detected in cells treated with Poly(I:C), while miR-6001-3p and miR-996-3p were exclusive to ABPV-infected cells. Using AmE-711 honey bee cells to study infections by different virus families could help further characterize these intra- and interspecific responses.

Previous studies have linked miR-6001-3p to Wnt, Hippo, and Notch signaling during the development of the larval gut in Asian honey bees (*A. cerana*). The Wnt signaling pathway has also been demonstrated to strongly inhibit the replication of SARS-CoV-2 and other RNA viruses *in vitro* while also reducing viral load, inflammation, and clinical symptoms in a murine model of COVID-19 infection. Given this, further studies that inhibit the expression of miR-6001-3p during viral infection could firm the linkage between Wnt signaling and viral pathogenesis.

### Limitations of the Study

As previously reported (Carillo-Tripp et al., 2016), the AmE-711 honey bee cell line is persistently infected with DWV. Persistent DWV infection in AmE-711 cells represents a confounding factor that could enhance or suppress miRNA expression caused by co-infection with another virus like ABPV. Ideally, studying ABPV-induced miRNA regulation would benefit from a virus-free honey bee cell line (Guo et al., 2020). To achieve this, approaches such as antiviral drug treatment, sub-cloning, and CRISPR/Cas13-mediated viral targeting could be used to eliminate DWV infection from AmE-711. Furthermore, combining honey bee cell lines with virus replicons. infectious clones, and CRISPR/Cas9-mediated genome editing could create a powerful platform for investigating molecular mechanisms of virus-host interactions. This would advance our understanding of viral pathogenesis, honey bee antiviral immunity, and more generally, the invertebrate immune response to viral infections.

## Data availability statement

The raw small RNA sequences were deposited into the NCBI Sequence Read Archive (SRA) repository under the BioProject ID PRJNA1173229.

## Author contributions

Conceptualization: DK, MG, JA, SK; Methodology: DK, MG; Investigation: DK, MG, JA, SK; Formal Analysis: DK, SK; Writing-Original Draft: DK, MG, JA, SK; Writing-Review & Editing: DK, MG, JA, SK; Funding Acquisition: MG, JA, SK; Supervision: SK

## Acknowledgment

The authors acknowledge Magnolia HPC at The University of Southern Mississippi supported by the National Science Foundation under the Major Research Instrumentation (MRI) program via Grant # ACI 1626217.

